# Enhanced Environmental Detection of Valley Fever Hotspots Near the US-Mexico Border Using Droplet Digital PCR

**DOI:** 10.1101/2025.08.10.669480

**Authors:** Jessica Paulette Segovia-Mota, Ricardo Eaton-González, Jimena Carrillo-Tripp, Meritxell Riquelme

## Abstract

Coccidioidomycosis (Valley Fever) is a reemerging, neglected fungal disease endemic to arid and semi-arid regions of the Americas, caused by the soil-dwelling fungi *Coccidioides* spp. Environmental detection remains challenging due to spatial heterogeneity, seasonal variability, low DNA abundance, PCR inhibitors, and lack of standardized methods. We conducted environmental surveillance in Baja California, Mexico, an understudied region near the U.S.-Mexico border, by collecting 74 soil samples from active rodent burrows across five locations. We evaluated droplet digital PCR (ddPCR) for *Coccidioides* detection and compared its performance to a nested PCR assay targeting the ITS1 region. ddPCR demonstrated greater sensitivity, detecting *Coccidioides* DNA at all sampling sites, whereas nested PCR detected it at only one. These findings highlight ddPCR as a sensitive and reliable method for environmental detection of *Coccidioides*. Integrating high-resolution molecular diagnostics with ecological data may enhance species distribution models and guide targeted public health interventions in endemic regions.

## Introduction

Coccidioidomycosis (CM), commonly known as Valley Fever, has been increasingly reported across the American continent, pointing to its potential reemergence as a public health concern (1). This disease is caused by two species of ascomycetous fungi belonging to the genus *Coccidioides*: *C. immitis* and *C. posadasii* (2). *Coccidioides immitis* is primarily found in California, in the United States, whereas *C. posadasii* prevails in southern Arizona, New Mexico, Texas, parts of Central America and South America (2–4), as well as various Mexican states, including Sonora, Nuevo León, Coahuila, and Baja California. Today, CM is considered one of North America’s most important endemic mycoses (5–7).

Despite high incidence rates of CM in the United States (estimated between 150,000 and 350,000 cases annually) (11), environmental detection of *Coccidioides* spp. remains limited. Positive isolates from soil samples are rare compared to clinical samples. Several factors contribute to this: non-targeted sampling strategies, limited understanding of the fungus’ ecological niche, presence of other fungi and bacteria that interfere with the isolation, and the inherent complexity and unpredictability of traditional culture-based isolation methods (8–11). Most attempts to isolate *Coccidioides* spp. from the environment have produced low yields. Even animal inoculation methods using soil-derived suspensions have shown only limited success (7, 9).

Similarly, attempts to detect *Coccidioides* DNA directly from soil-without prior fungal isolation-have given mixed results (12). In one study, 90 soil samples were collected from active rodent burrows in two semi-arid regions of Baja California, Mexico: Valle de las Palmas (VDP) and San José de la Zorra (SJZ). These samples were analyzed using nested PCR followed by diagnostic PCR targeting the ITS2 region of *Coccidioides*. While 32 samples (35.5%) tested positive −27 from VDP and five from SJZ-13.33% were later confirmed as false positives. This lack of specificity was attributed to high sequence similarity at the 5’ and 3’ ends of the ITS2 region among various fungal species, which led to amplification of non-target DNA from fungi such as *Aphanoascus canadensis, A. keratinophilus, A. verrucosus, A. pinarensis*, and *Penicillium* sp., in addition to *Coccidioides* spp. (12–14).

In a follow-up study, Vargas-Gastélum and colleagues (14) assessed fungal diversity in rodent burrows and surface soils during both winter and summer, using pyrosequencing and nested PCR. To reduce false positives, they designed primers targeting the ITS1 region, which offered greater specificity to *Coccidioides* spp. While pyrosequencing did not detect the pathogen, nested PCR identified *Coccidioides* DNA in 11 of 40 samples (27.5%), nine from burrows and two from surface soils.

Droplet digital PCR (ddPCR) is an emerging technique that significantly enhances detection sensitivity. It utilizes an emulsion of water and oil to generate approximately 20,000 droplets, each acting as an individual reaction site for PCR amplification (Droplet digital PCR, ddPCR, Technology, 2024). Each droplet may contain zero or one copy of the target sequence, allowing for highly accurate quantification (15). The ddPCR technique has already shown promise in detecting and quantifying a range of microorganisms, including fungi. For instance, when comparing qPCR and ddPCR for detecting *Aspergillus* species in the human respiratory tract (16), ddPCR identified 15 of the 16 samples as positive (93.8%), outperforming qPCR, which detected 13 (81.3%). Notably, ddPCR was more effective at detecting *A. terreus* at low concentrations (16). Another study comparing conventional end point PCR, real-time PCR, and ddPCR for detecting *Tilletia controversa* teliospores in soil samples found ddPCR to be the most sensitive, detecting down to with 2.1 copies/µL – making it roughly 100 times more sensitive than conventional PCR, and 3.8 times more sensitive than real-time PCR with 7.97 copies/µL (17).

Given these advantages, the aim of this study was to assess ddPCR’s ability to detect *Coccidioides* spp. in soil samples from sites previously identified as positive. Additionally, we aimed to screen other unexplored potentially endemic areas of Baja California, Mexico, flagged as hotspots for *Coccidioides* in recent species distribution models (18).

## Methods

### Sampling sites

Five sampling sites within two localities, VDP and SJZ, were selected based on previous reports confirming the presence of *Coccidioides* (12, 18). Locality VDP, located approximately 10 km southeast of Tijuana, Baja California, includes three sampling sites: Rancho Gilbert (RG), Rancho Las Golondrinas (RLG), and Rancho Carrizo (RC). Locality SJZ, situated about 32 km north of Ensenada, Baja California, comprises the Community of San José de la Zorra (CSJZ) and Agua Escondida (AE) (Figure 1).

**Figure 1.**
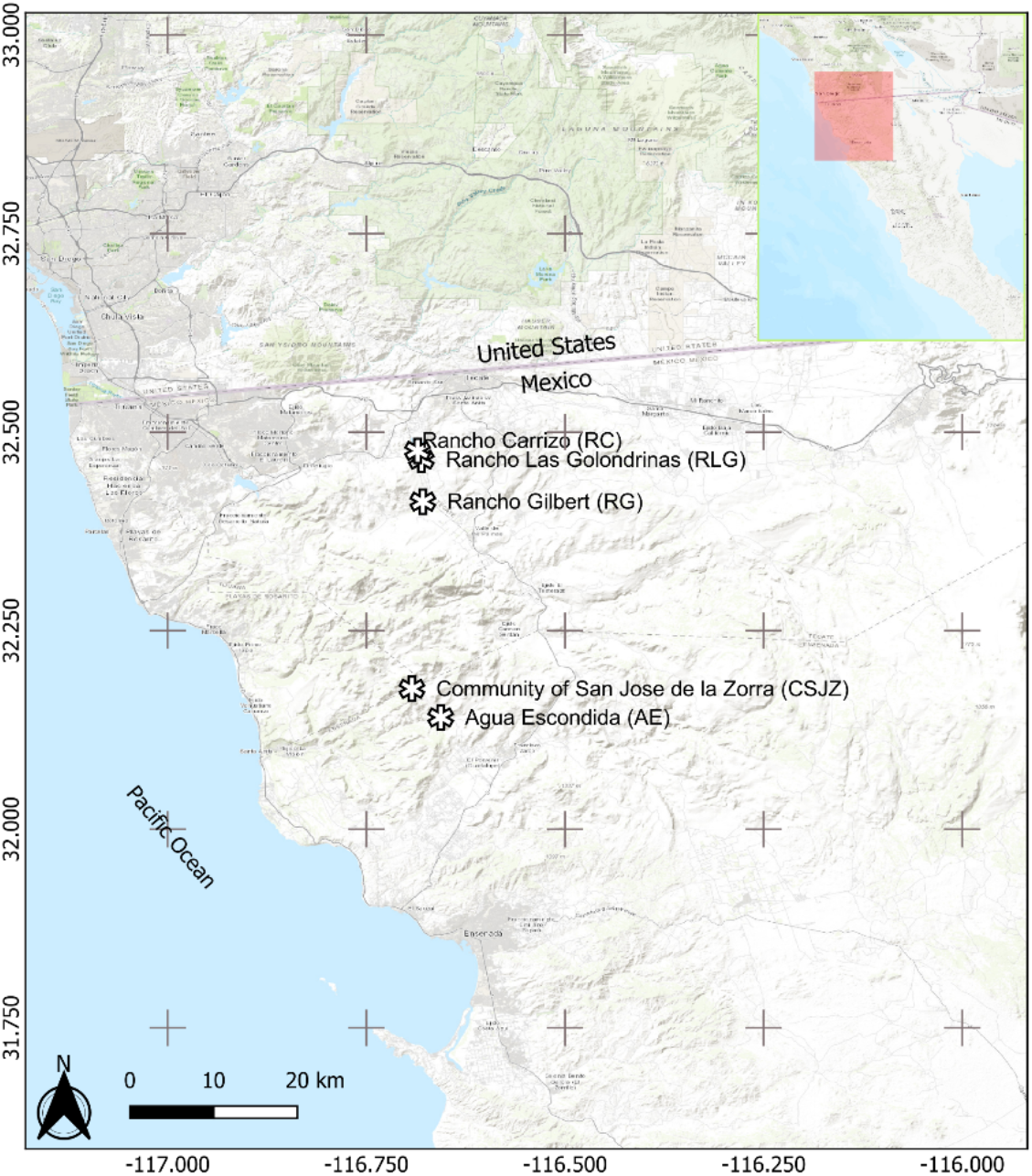
Location of the sampling sites in Baja California, Mexico: Rancho Gilbert (RG), Rancho Las Golondrinas (RLG), Rancho Carrizo (RC), Community of San José de la Zorra (CSJZ), and Agua Escondida (AE)

A total of 76 soil samples were collected: 20 from RG, 12 from RLG, 18 from RC, 14 from CSJZ, and 12 from AE (Supplementary Table 1). Sampling was directed to active rodent burrows, as previous studies have shown higher *Coccidioides* prevalence in burrows soil compared to surface soil (12-14, 19). Rodents are thought to create favorable microenvironments for fungal growth, providing warmth and moisture, without presenting disease symptoms, potentially acting as reservoirs (2, 13, 20).

Active burrows were identified by the absence of cobwebs or vegetation obstructing the entrance and the presence of loose, uncompacted soil, indicating recent use. Approximately 40 g of soil were collected from 10 to 15 cm inside the burrow entrance into sterile plastic jars. To avoid cross-contamination, the sampling spoon used was rinsed with chlorine between sites. As preventive safety measures, sterile gloves and face masks were used during sample handling.

At each site, data were recorded using a standardized form, capturing information such as GPS coordinates, temperature, wind speed, predominant vegetation (e. g., scrub, chaparral, forest, grassland, unvegetated/bare soil, or riparian), characteristics of the sampling site (e. g., burrow associated with a rock or plant, or without association), soil texture (coarse, medium, fine), soil compaction (compact or uncompacted), and moisture condition (dry, wet, mixed). Panoramic site photographs and images of the specific burrow and collected sample were also taken.

### DNA extraction from soil samples

DNA was extracted using the QIAGEN DNeasy PowerSoil Pro kit, following the manufacturer’s protocol with 250 mg of soil per extraction. DNA concentration and purity were assessed using a Thermo Scientific Nanodrop LITE spectrophotometer (A260, A280), with 1 μL of sample. DNA integrity was verified by electrophoresis on a 1% agarose gel. All DNA samples were then normalized to a final concentration of 10 ng/μL.

### Detection of *Coccidioides* spp. by nested PCR

Nested PCR was used to detect *Coccidioides* from the DNA extracted from the soil samples. The first PCR targeted the ITS1-5.8s-ITS2 region (approximately 900 bp) using primers NS1 and NLB4 (12, 21). Genomic DNA from *C. posadasii* (courtesy of Dr. Bridget Barker from the Institute of Pathogens and Microbiomes, Northern Arizona University, USA) was used as a positive control, while HPLC-grade water was used as a negative control. Amplified products were confirmed by electrophoresis on a 1% agarose gel. For the second round of amplification, a 1:10 dilution of the first PCR product was used. This reaction targeted the ITS1 region (approximately 120 bp), specific for *Coccidioides* spp. using primers ITS1CF and ITS1CR (14). The products were analyzed by electrophoresis on a 2% agarose gel. Positive amplicons were purified using a Qiaquick Gel Extraction Kit (Qiagen) and sequenced (Eton Bioscience Inc., San Diego, CA). Sequencing results were manually curated using MEGA (Molecular Evolutionary Genetics Analysis, version 11.0.13), and species identity was confirmed via BLASTn searches against the NCBI nucleotide database.

### Detection of *Coccidioides* spp. by ddPCR

Droplet digital PCR was performed on a QX200 Droplet Digital PCR system (Bio-Rad) after optimizing primer concentrations and annealing temperatures. The same primers used in the second nested PCR were adapted for ddPCR following Bio-Rad guidelines (2018), which recommend avoiding a G at the 5’ end, maintaining a GC content of 50-60%, ending the 3’ end with a G or C, and avoiding primer dimer formation. Each 21 μL reaction contained: 10.5 μL of QX200 ddPCR EvaGreen Supermix (2X, BioRad), 0.126 μL (60 nM) each of forward (ITS1CF-EVA 5’-AAGTGGCGTCCGGCTGCGCACCTCCCCCGCGG-3’) and reverse (ITS1CR-EVA 5’-ACGCCGCGCAAGGCGGGCGATCCCCGGC-3’) primers, 9.248 μL of Milli-Q grade water, and 1 μL (10 ng) of template DNA. Two types of negative controls were included: (1) no template controls (NTC, using HPLC-grade water), and (2) *Aspergillus fischeri* DNA (28.8 ng/μL, kindly provided by D. Martínez-Soto, CICESE). *Coccidioides posadasii* DNA (4.8 ng/μL) was used as a positive control. Droplets were generated using a QX200 Droplet Generator (Bio-Rad), transferred to a 96-well plate, sealed with aluminum foil (PX1 PCR Plate Sealer, Bio-Rad), and processed in a C1000 Touch Thermal Cycler (Bio-Rad). The amplification program consisted of an initial cycle of enzyme activation at 95°C for 10 min, followed by 40 cycles of: denaturation at 95°C for 30 s, annealing/extension at 70°C for 60 s, and stabilization at 4°C for 5 min, followed by inactivation at 90°C for 5 min. Following amplification, the plate was read using the QX200 Droplet Reader (Bio-Rad). Droplet analysis was performed using the QuantaSoft and QuantaSoft Analysis Pro software (Bio-Rad). The threshold to differentiate positive from negative droplets was manually set based on negative controls. The concentration of *Coccidioides* DNA in each sample was reported in copies/μL, calculated from the fraction of positive droplets.

## Results

### Yield and purity of DNA recovered from soil samples

The average A260/280 ratio across all samples was 1.69, with DNA purity consistently below While lower than ideal, these values are acceptable for downstream molecular applications, such as PCR and ddPCR.

### Detection of *Coccidioides* spp. by nested PCR

Out of the 76 total soil samples analyzed, only five samples-representing 25% of those collected from RG-tested positive for *Coccidioides* spp. after the second round of nested PCR. These positive samples (RG1, RG9, RG10, RG19, and RG20) exhibited amplicons within the expected size range (around 120 bp), falling between 100 and 200 bp (Figure 2). Sequencing and BLAST analysis of the five positive amplicons confirmed that all corresponded to *C. immitis*. In addition, sequencing of the positive control used throughout the PCR experiments verified it as *C. posadasii*, as expected (Supplementary Table 2).

**Figure 2.**
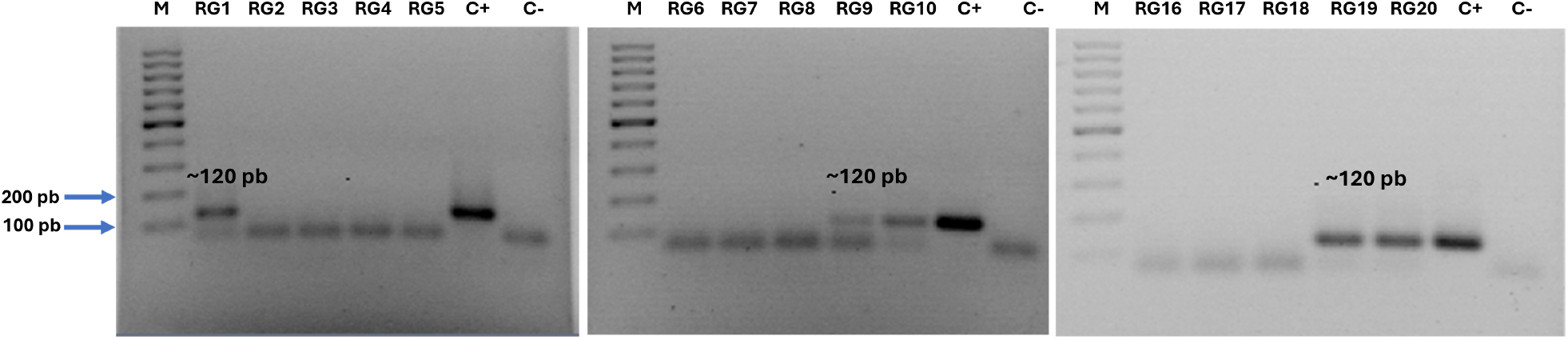
Nested PCR amplification of the ITS1 region of 20 samples from Rancho Gilbert. Agarose gel electrophoresis (2%) stained with ethidium bromide. M: GeneRuler™ ladder (100 bp), RG1-RG20: samples, C+: positive control (*C. posadasii* DNA), C-: negative control (HPLC water).

### Detection of *Coccidioides* spp. by ddPCR

For all ddPCR reactions, over 10,000 droplets were successfully generated per sample, ensuring sufficient sensitivity and robustness. Among the negative controls used for the RG samples, two of the three NTC and both *Aspergillus fischeri* DNA controls showed no positive droplets. However, one NTC (NTC1) produced a single droplet above the fluorescence threshold (0.0663 copies/µL; Supplementary Table 3), likely due to minor contamination during sample handling. Similarly, isolated single-droplet signals were observed in two of the three NTCs for the other sample sets (RC, RLG, CSJZ, and AE). Based on this, samples with only one positive droplet were conservatively classified as negative.

Of the 20 samples from RG, 16 showed amplification with multiple positive droplets, indicating the presence of *Coccidioides* DNA (Figure 3A). The five samples previously identified as positive by nested PCR (RG1, RG9, RG10, RG19, and RG20) were all confirmed by ddPCR. In addition, 11 other samples also tested positive by ddPCR, excluding RG4, RG8, RG11, and RG12, which remained negative. Collectively, the ddPCR indicated a detection rate of 80% for RG. Among the 18 samples from RC, which were all negative by nested PCR, five samples (RC1, RC5, RC8, RC10, and RC18) showed positive droplet amplification by ddPCR, yielding a positivity rate of 27.7% (Figure 3B). For RLG, only one of the 12 samples (RLG4) tested positive by ddPCR (Figure 3C), resulting in an 8.3% positivity rate.

**Figure 3.**
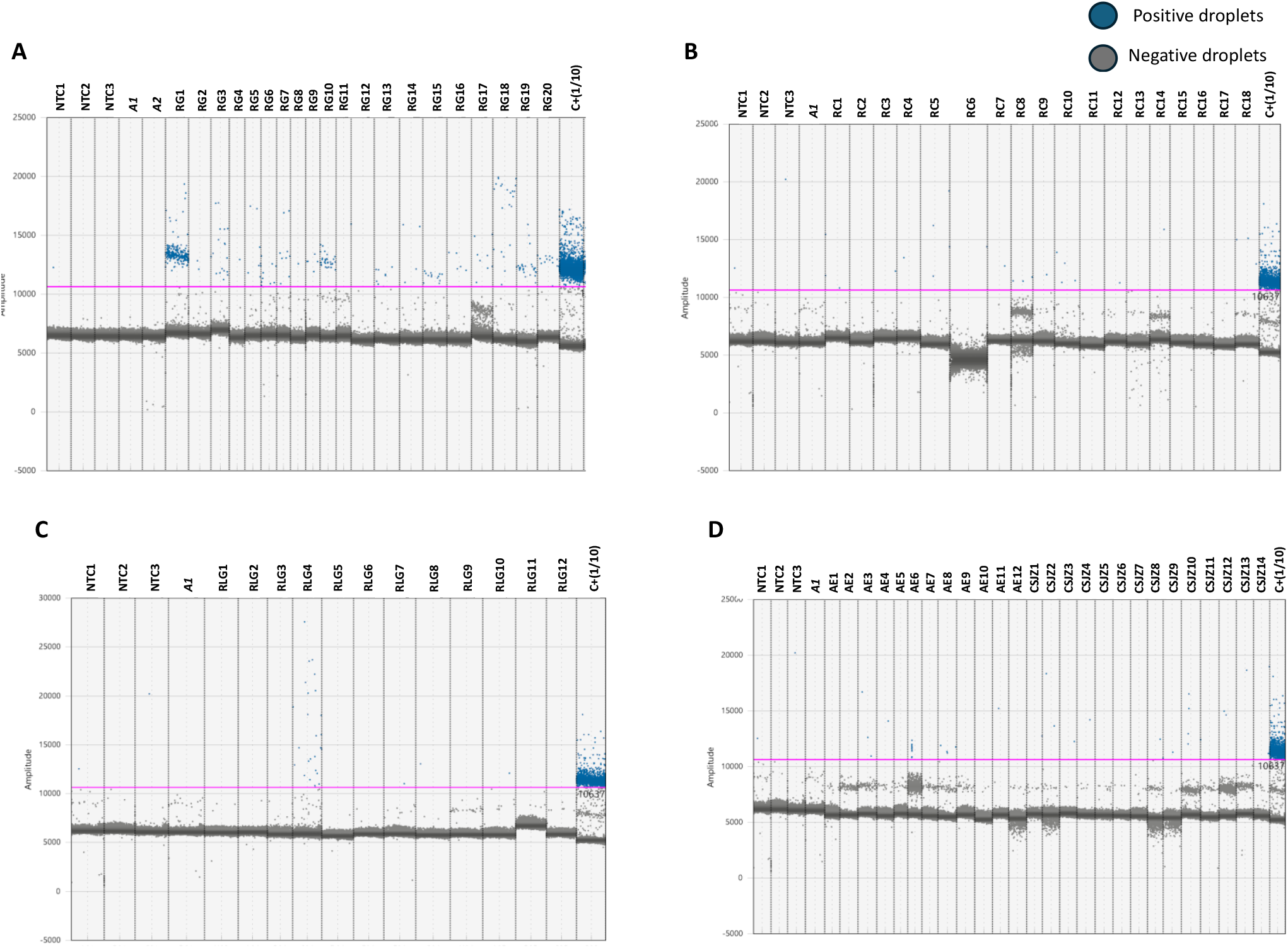
Droplet distribution diagram of soil samples tested by ddPCR. A) Rancho Gilbert (RG), B) Rancho Carrizo (RC), C) Rancho las Golondrinas (RLG), D) Agua Escondida (AE), and Community of San José de la Zorra (CSJZ). Blue dots correspond to positive droplets; gray dots correspond to negative droplets. *A1 and A2: Aspergillus fischeri*; C+: *C. posadasii* (4.8 ng/μL); NTC1, NTC2, and NTC3 represent negative controls (HPLC water), and the pink line represents the threshold.

For the other locality comprising CSJZ and AE, none of the 26 samples tested positive by nested PCR. However, ddPCR detected *Coccidioides* DNA in eight of these samples (AE3, AE4, AE6, AE8, CSJZ2, CSJZ8, CSJZ10, and CSJZ12), resulting in an overall positivity rate of 30.7% for this locality (Figure 3D; Figure 4).

**Figure 4.**
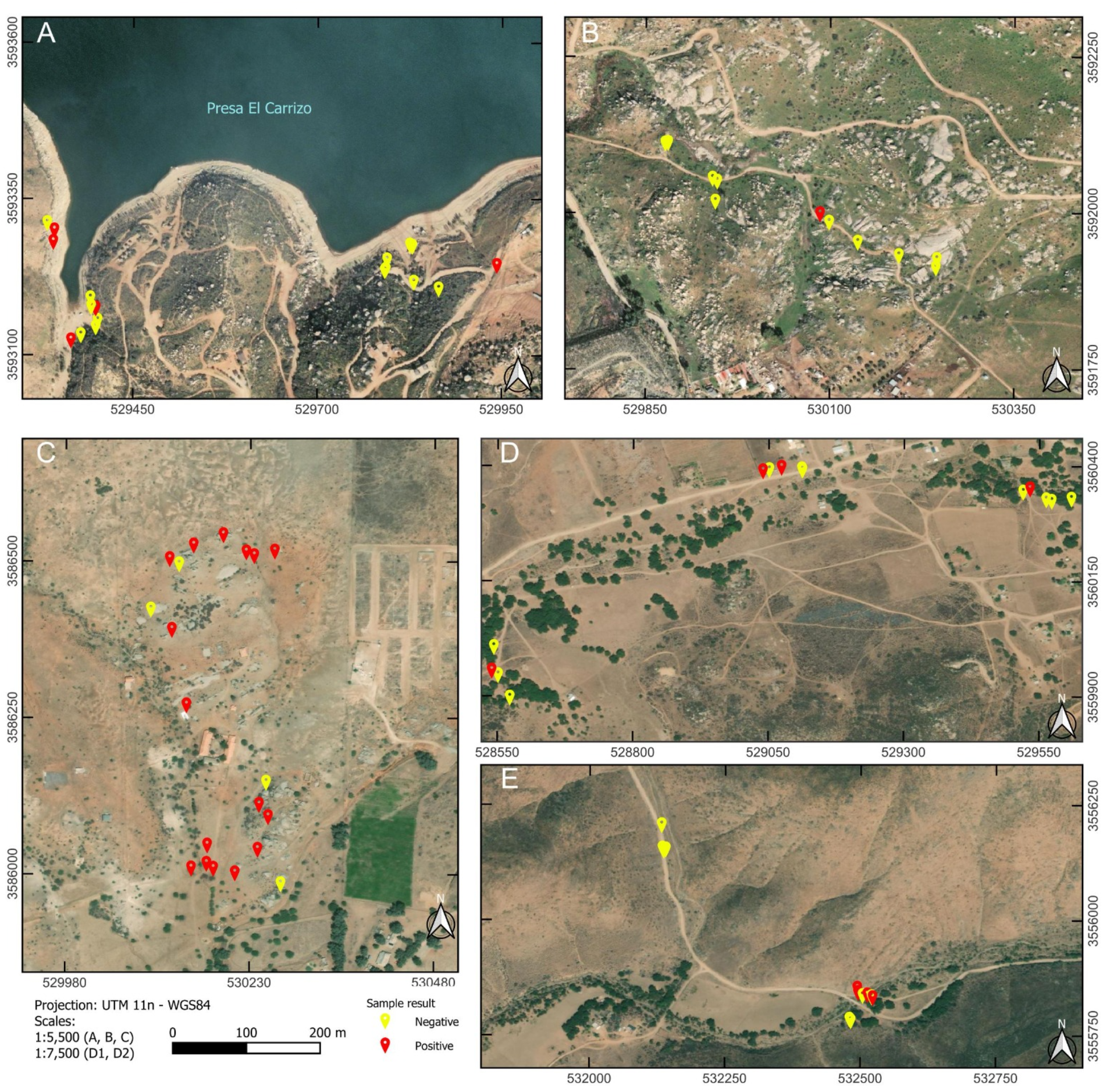
Negative and positive sites by ddPCR. A) Rancho Carrizo (RC), B) Rancho las Golondrinas (RLG), C) Rancho Gilbert (RG), D) Community of San José de la Zorra (CSJZ), and E) Agua Escondida (AE).

## Discussion

This research demonstrates that ddPCR is a more sensitive and effective technique than nested PCR for detecting *Coccidioides* spp. in environmental soil samples (Table 1). Our results show that ddPCR not only confirmed all positive samples previously identified by nested PCR in the VDP locality, but also detected a significantly higher number of positive samples in both VDP and SJZ, including areas previously considered negative. This study highlights ddPCR’s capacity to uncover cryptic environmental reservoirs of *Coccidioides* and potentially other pathogens that may contribute to human exposure and disease.

**Table 1.** Results obtained by Nested PCR and ddPCR.

| Samples (76) | Nested PCR | ddPCR |
| --- | --- | --- |
| 20 RG | 5 | 16 |
| 18 RC | 0 | 5 |
| 12 RLG | 0 | 1 |
| 14 CSJZ | 0 | 4 |
| 12 AE | 0 | 4 |

The ability of ddPCR to detect low quantities of *Coccidioides* DNA allowed us to expand the known distribution of this pathogen within the two localities studied – VDP and SJZ. While nested PCR detected *Coccidioides* in only five (all from VDP) of 76 total samples, ddPCR confirmed those same five samples and additionally detected *Coccidioides* DNA in 17 of 50 VDP samples and 8 of 26 SJZ samples. These findings suggest that ddPCR can reveal a more accurate and complete environmental presence of *Coccidioides*, even in areas where detection has historically been challenging.

In earlier studies, nested PCR using primers targeting the ITS2 region reported positivity rates of 74% for VDP and 10% for SJZ, with burrow samples contributing significantly to the detection (6, 13). Although all samples were taken from active burrows in our study, we observed a lower detection rate via nested PCR: 25% for VDP and 0% for SJZ. The discrepancy could be attributed to smaller sample sizes, temporal environmental variability, or differences in primer sets and target regions (Supplementary Table 5). These discrepancies reinforce the importance of selecting highly sensitive detection methods, such as ddPCR, for pathogen surveillance in heterogeneous environments and for cryptic human and environmental pathogens, such as *Coccidioides*.

It is noteworthy that earlier studies employed ITS2F and ITS2R primers (6, 13), while our nested PCR protocol used the ITS1CF and ITS1CR primers developed by Vargas-Gastélum et al. (14). Their approach successfully detected 11 positive samples (9 from burrows) out of 40, suggesting improved specificity for *Coccidioides* spp.

The technique of ddPCR has previously demonstrated high sensitivity in clinical settings for detecting fungal pathogens such as *Pneumocystis, Aspergillus*, and *Cryptococcus* in immunocompromised patients (22). Our study marks the first application of ddPCR for detecting *Coccidioides* spp. in environmental soil samples, demonstrating its potential as a scalable and precise tool for environmental surveillance and early warning systems in endemic regions of this pathogen and others of clinical importance.

Although ddPCR results showed clear positives at each site, threshold determination is critical when working with environmental DNA. We manually set a high fluorescence threshold to reduce false positives, excluding isolated droplets observed in negative controls. However, due to the complexity of soil matrices and the possibility of droplet misclassification or background fluorescence, sometimes referred to as the “significant rainfall” effect, some degree of false positives cannot be completely ruled out. This limitation is especially relevant since ddPCR does not permit sequencing of the amplified products, making it difficult to distinguish true positives from potential artifacts (23). To address this problem, we adopted a cautious interpretative approach, excluding samples with only a single droplet above the threshold. While this minimizes overestimation, it also raises the possibility of underreporting true positives. Nevertheless, these borderline cases may still represent genuine low-copy *Coccidioides* DNA, emphasizing the need for additional validation studies to refine ddPCR thresholds and improve specificity for environmental applications.

From a public health perspective, there is urgent need to improve environmental detection of *Coccidioides* amid increasing Valley Fever incidence and expanding distribution, possibly linked to climate change and land-use transformations (18, 24, 25). This work represents a foundational effort to apply ddPCR for the environmental detection of *Coccidioides* spp. The results highlight the method’s superior sensitivity over nested PCR and its potential for surveillance based on environmental samples (soil samples), particularly in vulnerable regions with similar ecological characteristics, to uncover new endemic areas previously undetected by conventional techniques. The enhanced sensitivity of ddPCR offers opportunities to improve spatial risk mapping, anticipate outbreak potential, and strengthen early detection strategies. Further research using larger sample sets, additional molecular markers, and sequencing confirmation will be essential to improve detection accuracy and our understanding of *Coccidioides* ecology in soil environments, supporting more informed public health interventions across endemic and emerging regions.

## Acknowledgements

We are grateful to the Consejo Nacional de Humanidades, Ciencias y Tecnologías (CONAHCYT) of Mexico for providing a scholarship to PSM. We would like to thank Ofelia Sastré and Jonatan Aguilar from UABC for technical help during the 2023 sampling and Ofelia Candolfi for providing the DNeasy PowerSoil ProKit from QIAGEN. We thank Dolores Camacho for technical assistance with nested PCR, and Karen García, Idalia Montesinos, and Sócrates Avilés (Biorad) for technical assistance with ddPCR.

## Statement

During the preparation of this work the authors declare the use of ChatGPT to check grammar and improve readability and language of the work. After using this tool, the authors reviewed and edited the content as needed and take full responsibility for the content of the published article.

## Supplementary Material

ST1. Coordinates of the sampling sites. See separate ST1 Excel file.

ST2. Fungal species identified by sequencing of the amplicons of the ITS1 region after extraction of environmental DNA from five soil samples.

| Sample | Fungal species of amplicons by soil DNA extraction (ITS1 region) | E value | % Identity | Query Cover |
| --- | --- | --- | --- | --- |
| RG1 | *Coccidioides immitis* | 3e-38 | 98.10% | 95% |
| RG9 | *Coccidioides immitis* | 1e-33 | 98.88% | 89% |
| RG10 | *Coccidioides immitis* | 2e-36 | 100% | 88% |
| RG19 | *Coccidioides immitis* | 2e-35 | 98.91% | 85% |
| RG20 | *Coccidioides immitis* | 3e-31 | 97.59% | 81% |
| + Control (*C. posadasii*) | *Coccidioides posadasii* | 6e-40 | 98.06% | 91% |

ST3. Table of data of the results obtained from soil samples by ddPCR. See separate ST3 Excel file.

ST4. Dates of samplings carried out by different authors.

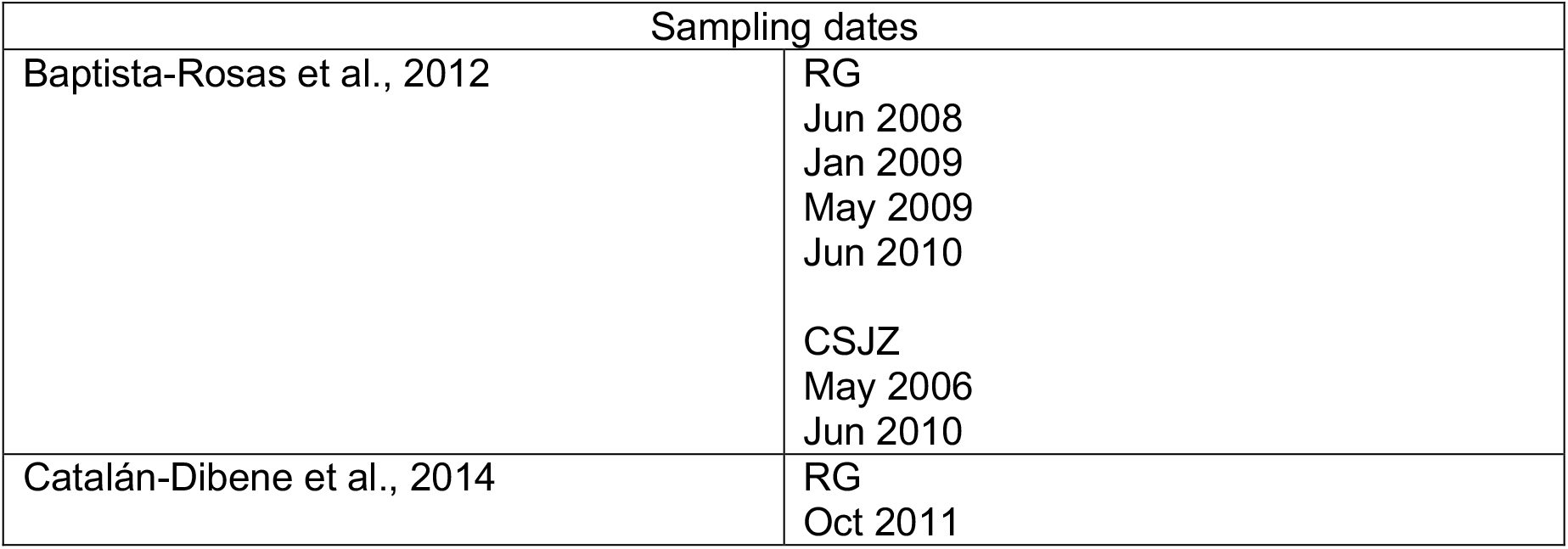

ST3. Table of data of the results obtained from soil samples by ddPCR. See separate ST3 Excel file. ST4. Dates of samplings carried out by different authors.
| Sampling dates |  |
| --- | --- |
| Baptista-Rosas et al., 2012 | RG<br>Jun 2008<br>Jan 2009<br>May 2009<br>Jun 2010 |
|  | CSJZ<br>May 2006<br>Jun 2010 |
| Catalán-Dibene et al., 2014 | RG<br>Oct 2011 |

**First author biography**

Jessica Paulette Segovia-Mota is a biologist with a Master’s degree in Life Sciences. She has a deep interest in pathogenic fungi, dedicating much of her professional and academic pursuits to understanding their role in ecosystems and their potential impact on human health.

